# Single-cell transcriptomic analysis of mouse liver reveals nonparenchymal cells’ intricate responses to PCB126 exposure

**DOI:** 10.1101/2023.02.13.528414

**Authors:** Fengjun Xu, Yulong Fu, Jiaxuan Yang, Chunna Yu, Chaofeng Shen

**Affiliations:** Department of Chemical and Environmental Engineering, University of California, Riverside, California 92521, United States; Department of Environmental Engineering, College of Environmental and Resource Sciences, Zhejiang University, Hangzhou 310058, P. R. China; HanGene Biotech, Xiaoshan Innovation Polis, Hangzhou 311200, P. R. China; College of Life and Environmental Sciences, Hangzhou Normal University, Hangzhou 311121, P. R. China; Zhejiang Provincial Key Laboratory for Water Pollution Control and Environmental Safety, Hangzhou 310058, P. R. China

**Keywords:** PCB126, PCB toxicity, single-cell RNA sequencing, aryl hydrocarbon receptor, immune response

## Abstract

Polychlorinated biphenyls (PCBs) are ubiquitous and representative pollutants that pose great health risks. While cells’ responses to dioxin-like PCBs tend to be studied on a bulk scale, few studies have been made from a single-cell level. Here, by using single-cell RNA sequencing, we depicted a detailed landscape of hepatic nonparenchymal cells’ intricate responses to PCB126 exposure. A total of 13 clusters were identified. Notably, PCB126 exposure resulted in cell-type-specific gene expression profiles and genetic pathways. By analyzing genes related to aryl hydrocarbon receptors, we discovered that PCB126 induced the canonical genomic AhR pathway mainly in endothelial cells. In contrast, other cell types showed little induction. Enrichment pathway analysis indicated that immune cells changed their transcriptional patterns in response to PCB126. ScRNA-seq is a powerful tool to dissect underlying mechanisms of chemical toxicity regarding biological heterogeneity. Taken together, our study not only extends our current understanding of PCB126 toxicity, but also emphasizes the importance of *in vivo* cell heterogeneity in environmental toxicology.

## Introduction

Polychlorinated biphenyls (PCBs), which were industrially manufactured in a large amount and earned various applications in the 20^th^ century, now have been recognized to pose great health risks (Safe 1994; Zhu et al. 2022). Although their production has already been banned all over the world, there were about 1.3 million tons of legacy PCBs produced from 1930 to 1993 (Breivik et al. 2002). In addition to this legacy, there are still PCBs inadvertently produced (Liu et al. 2020). Due to their high stability and recalcitrance in the environment, PCBs recirculate in air, water and soil, gradually bioaccumulating in upper animals through food chains (Xia et al. 2012). In terms of biological effects, PCBs could cause a series of negative immunological, neurological and endocrinological consequences (Crinnion 2011). Many studies discovered that PCB exposure is highly correlated with various diseases, like cancer (Zani et al. 2017), diabetes (Shi et al. 2019) and nonalcoholic fatty liver disease (Armstrong and Guo 2019; Wahlang et al. 2019; Jin et al. 2020; Shan et al. 2020).

Based on the structural and toxicity similarities to 2,3,7,8-tetrachlorodibenzo-p-dioxins (TCDDs), PCBs are classified into two categories–dioxin-like PCBs (DL PCBs) and non-dioxin-like PCBs (NDL PCBs). Compared with NDL PCBs, DL PCBs are more toxic, because they have the potent ability to bind to the aryl hydrocarbon receptor (AhR), which further activates the AhR pathway. When ligands like TCDDs and DL PCBs bind to AhR, the complex translocates into the nucleus, further binding to the aryl hydrocarbon receptor nuclear translocator (ARNT) to form a heterodimer. This heterodimer then modulates the expression of target genes by binding to xenobiotic responsive elements (XRE) and coregulators (Larigot et al. 2018). Target genes include *ahrr*, *cyp1a1*, *cyp1a2*, *cyp1b1*, etc. (Manikandan and Nagini 2017). The gene *ahrr* acts as the AhR repressor to form a negative regulatory loop (Baba et al. 2001), while *cyp1a1*, *cyp1a2* and *cyp1b1* belong to the cytochrome P450 (CYP) superfamily (Nebert and Dalton 2006). Regulations of these genes will facilitate the metabolism and excretion of xenobiotics. In this process, however, reactive oxygen species (ROS) are also generated and accumulated (Schlezinger et al. 1999), resulting in oxidative damage to intracellular substances and organelles. Among various kinds of DL PCBs, PCB126 (3,3’,4,4’,5-pentachlorobiphenyl) is thought to be the most potent and environmentally relevant one (Parvez et al. 2013). It was reported that up to 26% activation of AhR caused by environmental pollutants may be attributable to PCB126 (Shi et al. 2019).

As the largest solid organ in the body, the liver is critical for its metabolic and immune functions, including the detoxification of exogenous pollutants like PCBs. The liver is comprised of hepatocytes (parenchymal cells), which make up about two-thirds of its total cell population (60%-70%), and nonparenchymal cells (NPCs, 30%-40%). NPCs include endothelial cells (~50%), Kupffer cells (resident macrophages, ~20%), lymphocytes (~25%), cholangiocytes (~5%) and hepatic stellate cells (less than 1%) (Racanelli and Rehermann 2006). Hepatocytes contribute to a wide range of metabolic activity including xenobiotic metabolism (Washabau et al. 2012). Since hepatocytes are in charge of the major clearance work of DL PCBs, their responses have been extensively investigated (Zhang et al. 2012; Mesnage et al. 2018; Chen et al. 2020b). On the other hand, NPCs can be influenced as well. It was discovered that PCB126 is able to induce macrophage polarization and inflammation through AhR and NF-κB pathways (Wang et al. 2019). Besides, TCDDs can disrupt HSC activity and increase the activation of human HSCs (Harvey et al. 2016). While most studies focus on hepatocytes’ contributions, NPCs’ behaviors and roles in fighting against DL PCBs tend to be marginalized.

In recent years, single-cell RNA sequencing (scRNA-seq) has been brought into focus. It has the unparalleled ability to identify potential differences in gene expression of cells among different cell types and transcriptome disparity across cells of the same phenotype (Eberwine et al. 2014). Due to its powerful capability, scRNA-seq has flourished in bioscience, especially hepatology (Ramachandran et al. 2020). By using scRNA-seq, MacParland et al. depicted a map of the cellular landscape of human liver, and deepened our understanding of the human liver (MacParland et al. 2018). Meanwhile, scRNA-seq combined with single-molecule fluorescence *in situ* hybridization also helped complete a detailed genome-wide reconstruction of the spatial division of labor in mouse livers (Halpern et al. 2017). These studies all indicate the sophisticated structure of liver. Recently, this technique has also been applied to study the toxic effects of environmental pollutants like bisphenol A (Chen et al. 2020a), microplastics (Gu et al. 2020) and nanoplastics (Liu et al. 2021). ScRNA-seq empowers us to investigate toxic effects in a single-cell level, making the so-called “precision toxicology” possible (Zhang et al., 2017).

Although it has already been discovered that several hepatic NPCs can be influenced by DL PCBs, a detailed landscape remains to be investigated. Herein, by applying scRNA-seq, we aimed at showing a whole picture of hepatic NPCs’ intricate responses to PCB126 exposure in a mouse model. We found that both cells of different types and cells of the same type responded to PCB126 in a different manner. PCB126 induced the canonical AhR genomic pathway mainly in endothelial cells, and PCB126 gave rise to immune responses in multiple immune cells. This study not only profiles PCB126 toxicity from a single-cell level, but also provides insights into the application of scRNA-seq in environmental toxicology.

## Materials and Methods

### Animal husbandry and PCB exposure

All procedures related to animals were agreed upon and in accordance with the guideline of the Zhejiang University Animal Care and Use Committee. Male C57BL/6J mice (8 weeks old, ~20 g) were ordered from Shanghai SLAC Laboratory Animal Co., Ltd. (Shanghai, China). Before the experiment, mice were allowed to acclimatize for seven days. Two mice were involved in the experiment. Mice were given a common diet ordered by Slacom Co., Ltd. (Shanghai, China), and treated with either peanut oil (vehicle control) or PCB 126 (20 μg/kg/day, purchased from AccuStandard, Inc. (New Haven, CT)), respectively. The dose was determined by high-level human exposures in reality (Shi et al. 2019). PCB126 dissolved in isooctane (100 μg/mL) was added into peanut oil with a theoretical concentration of 4 μg/mL, and isooctane was removed by blowing a mild stream of nitrogen for 4 h. Each time 0.1 mL of peanut oil would be intragastrically administrated per 20 g weight to reach the concentration 20 μg/kg/day. Mice were housed with constant temperature (21°C-25°C) and humidity (45%-65%) in a light-controlled room (with light from 8:00 to 20:00), provided with food and water *ad libitum*. The exposure lasted for five weeks.

### Liver collection, dissociation and cell filtration

Mice were killed by cervical dislocation. The livers were carefully collected, rinsed and preserved in 1× PBS at 2°C to 8 °C, ready for downstream procedures. After the collection, livers were dissociated by using TissueLyser Type-TM Enzyme Blends (Jingxin Co., Ltd. (Shanghai, China)), under the manufacturer’s instructions. Centrifugation and filtration were performed to filter out hepatocytes, as previously described (Xiong et al. 2019), with minor modifications. In brief, the liver suspension was centrifuged at 300 g for 5 min several times and then passed through a 30 μm nylon cell strainer. Red blood cells were lysed by 0.8% NH_4_Cl treatment. The principle has been set to assure the quality of each sample that living cell numbers are larger than 10^6^ per milliliter, and that cell viability is at least 80%. These were confirmed via trypan blue staining and using a hemocytometer, respectively.

### Single-cell isolation, library preparation and sequencing

All procedures were processed based on the general protocols of 10x Genomics. Briefly, prior to loading onto the 10x Genomics single-cell-A chip, samples were diluted to an appropriate volume in which each sample was calculated for a targeted capture of 6,000 cells. After the droplet generation performed by single-cell microfluidic chips, samples were transferred onto a pre-chilled 96-well plate. cDNA libraries were prepared following the Single Cell 3’ Reagent Kits v3.1 user guide. Reverse transcription, cDNA purification and amplification were performed successively. Sequencing was conducted on the platform Illumina NovaSeq. 6000 with the help of HanGene Biotech Co., Ltd. (Hangzhou, China). All source data generated by single-cell RNA sequencing are uploaded to NCBI’s Gene Expression Omnibus (GEO), and easily accessible through the GEO series accession number (GSE190084).

### Data preprocessing

The software CellRanger v6.1.1 (10x Genomics, Pleasanton) and the package Seurat v4.0.1(Hao et al. 2021) in R language were applied to preprocess raw data. Raw sequence data (bcl files) were converted into fastq files and demultiplexed by the mkfastq command in CellRanger. The count command was used to perform the unique molecular identifiers and cell barcode deconvolution, as well as generate other important outputs (barcordes.tsv.gz, features.tsv.gz and matrix.tsv.gz) for downstream steps. Gene expression matrices were converted into Seurat objects using the CreateSeuratObject function. Quality was controlled by the subset function of Seurat in two steps. Firstly, all genes expressed in at least 10 cells and all cells with at least 200 detected genes were identified and included. Secondly, a minimum of 1500 and a maximum of 40000 were selected for unique gene count, and the percentage of mitochondrial genes > 0.5 was filtered out (MacParland et al. 2018). By using the NormalizeData function, filtered gene expression matrices were normalized by the total number of unique molecular identifiers per cell, multiplying by a scale factor (10000) and log-transformed. Top 1000 features were selected using the FindVariableFeatures function. The influence of cell circle and batch effects were taken into consideration by applying the CellCircleScoring function of Seurat and the package harmony v0.1.0 (Korsunsky et al. 2019), respectively.

### Cell clustering, differential expression and annotation

Cell clustering and differential expression were performed using Seurat. The RunPCA function was performed to run principal component analysis, and the RunTSNE function was used to perform t-distributed stochastic neighbor embedding (t-SNE) dimensionality reduction with the top 15 principal components, in which cells with similar expression signature genes and similar principal components localize near each other. The FindClusters function was used with a resolution of 0.2 to distinguish clusters. By using the FindNeighbors function, the clustering algorithm, K-nearest neighbor graph-based on the Euclidean distance in principal component analysis (PCA) space, was constructed to iteratively group similar cells together (Zhu et al. 2020).

The FindAllMarkers function of Seurat was used to identify marker genes of each cell cluster, and the FindMarkers function was conducted to identify differentially expressed genes (DEGs) between each cluster in the control group and PCB126 group. The identified clusters were firstly annotated by utilizing the package SingleR (Aran et al. 2019). The built-in data sets ImmGenData and MouseRNAseqData in the package celldex v1.2.0 (Aran et al. 2019) were chosen as references. To assure the accuracy of annotation, the gene marker list of each cluster was manually checked against known gene markers (Xiong et al. 2019; Zhu et al. 2020). The public database CellMarker (Zhang et al. 2019) and PanglaoDB (Franzén et al. 2019), which provide a plethora of gene markers based on previous single-cell research, were also utilized.

### GO and KEGG enrichment pathway analysis

Enrichment pathway analysis was performed on g:Profiler (Raudvere et al. 2019), a web server for functional enrichment analysis, with default parameters. The dataset containing DEGs was compared against the Gene Ontology (GO) and Kyoto Encyclopedia of Genes and Genomes (KEGG) reference gene sets. GO biological process (BP) enriched pathways and KEGG enriched pathways were identified. For immune cells, the interaction network of specific significant pathways was visualized by Enrichment Map v3.3.3 (Merico et al. 2010) in Cytoscape v3.8.2. Related pathways were grouped into a theme and labeled by AutoAnnotate v1.3.3 (Kucera et al. 2016) in Cytoscape v3.8.2.

### Statistical Analysis

When using the FindMarkers function of Seurat to find upregulated or downregulated genes, only those detected in a minimum of 10% cells in either group and having an average difference of at least 0.25 (log scale) between the two groups were tested. Statistical significance was calculated using the Wilcoxon rank-sum test, and p-value adjustment was performed using Bonferroni correction based on the total number of genes in the data set, which are embedded in Seurat (Zhu et al. 2020). An adjusted p-value < 0.05 was considered significant.

## Results

### Clustering and cell-type identification

After the filtration, a total of 4128 cells were sequenced and included, with 2066 in the control group and 2062 in the PCB126 group. Thirteen distinct cell clusters were identified (**Fig. 1a**). The liver is a complicated and sophisticated organ. In fact, cells of the same type can be different. This can be attributable to cell differentiation and their locations in the liver. With a resolution of 0.2, it was observed that endothelial cells and neutrophils were divided into several subpopulations (**Fig. 1a**). Endothelial cells were divided into Endo-1 (endothelial cells-1), Endo-2 and Endo-3, while neutrophils were separated as Neutro-1 (neutrophils-1), Neutro-2 and Neutro-3.

**Fig. 1.**
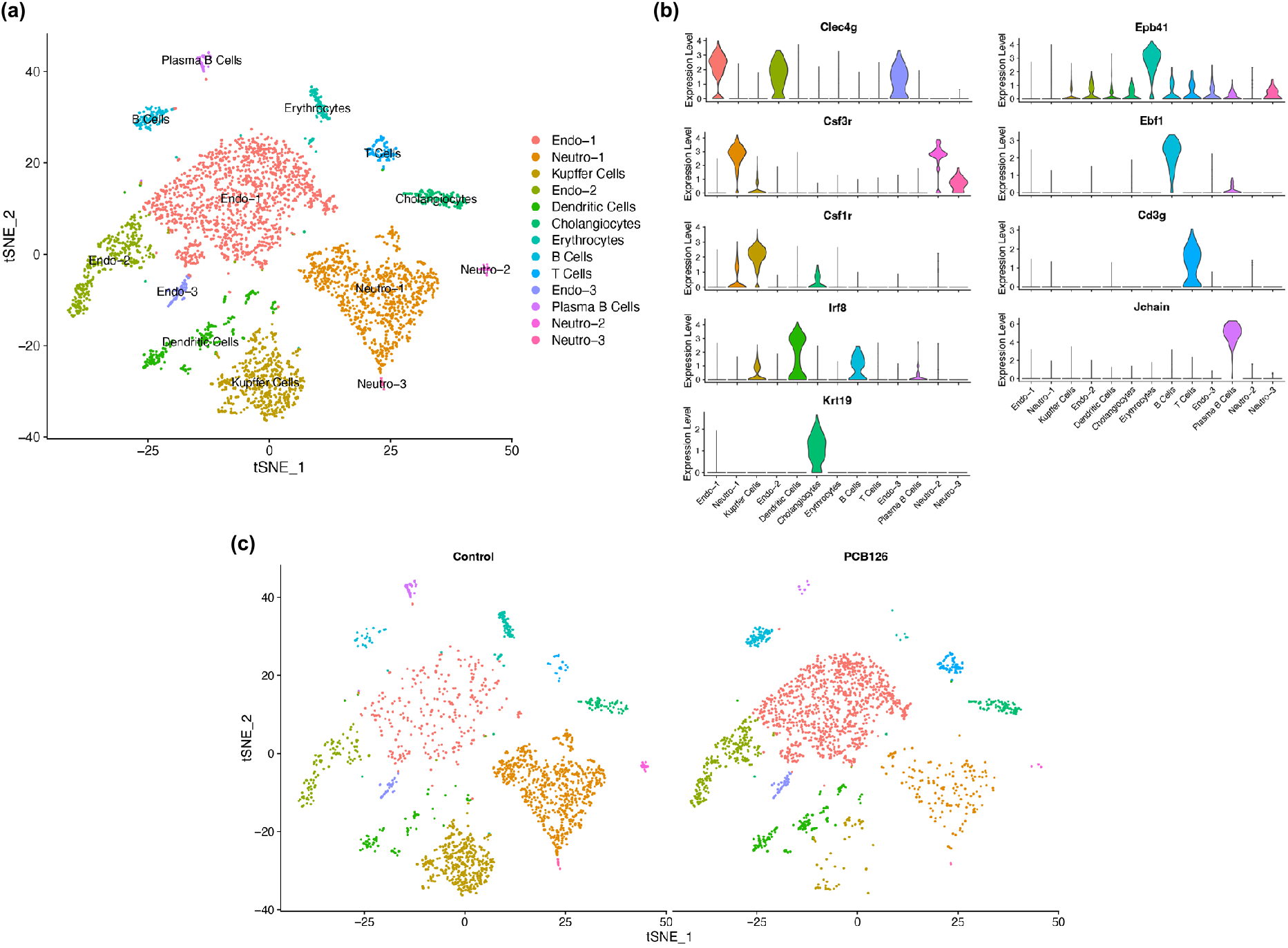
Clustering and cell-type identification using marker genes. **(a)** Clustering with annotation. **(b)** The expression of some marker genes in each cluster. **(c)** Cell distribution in two groups. Endo, endothelial cells; Neutro, neutrophils

All clusters are annotated according to known marker genes. Endothelial cells, Kupffer cells, neutrophils, dendritic cells, plasma B cells, B cells, T cells, cholangiocytes and erythrocytes were identified. Marker genes successfully separated different cell types apart (**Fig. 1b**). The gene *Clec4g* is a well-known marker gene for hepatic endothelial cells (Aizarani et al. 2019). Besides, *Ebf1* was important in the development of B cells (Zhang 2003), and *Cd3g* encodes the protein which is part of the T-cell receptor-CD3 complex (Charmley et al. 1989). Other distinguishing marker genes were also in agreement with the annotation. Cell types and their corresponding marker genes were listed in **Table S1**. The heatmap of the top 10 marker genes of each cluster were also plotted to help annotation (**Fig. S1**).

While all clusters appeared in two groups, their distributions were different (**Fig. 1c**). Compared with the control group, the PCB126 group had a larger proportion of endothelial cells and a smaller proportion of Kupffer cells.

### PCB126 results in cell-type-specific gene expression profiles and genetic pathways

Different DEG numbers of each cluster were obtained between the control group and the PCB126 group (**Fig. 2a**). Kupffer cells were the most altered cell type, with 696 DEGs. 261, 248 and 230 DEGs were detected in Endo-1, Neutro-1 and Endo-2, respectively. The rest clusters showed smaller DEG numbers. A total of 1595 DEGs was identified in all clusters, and different cell clusters occupied different proportions (**Fig. 2b**).

**Fig. 2.**
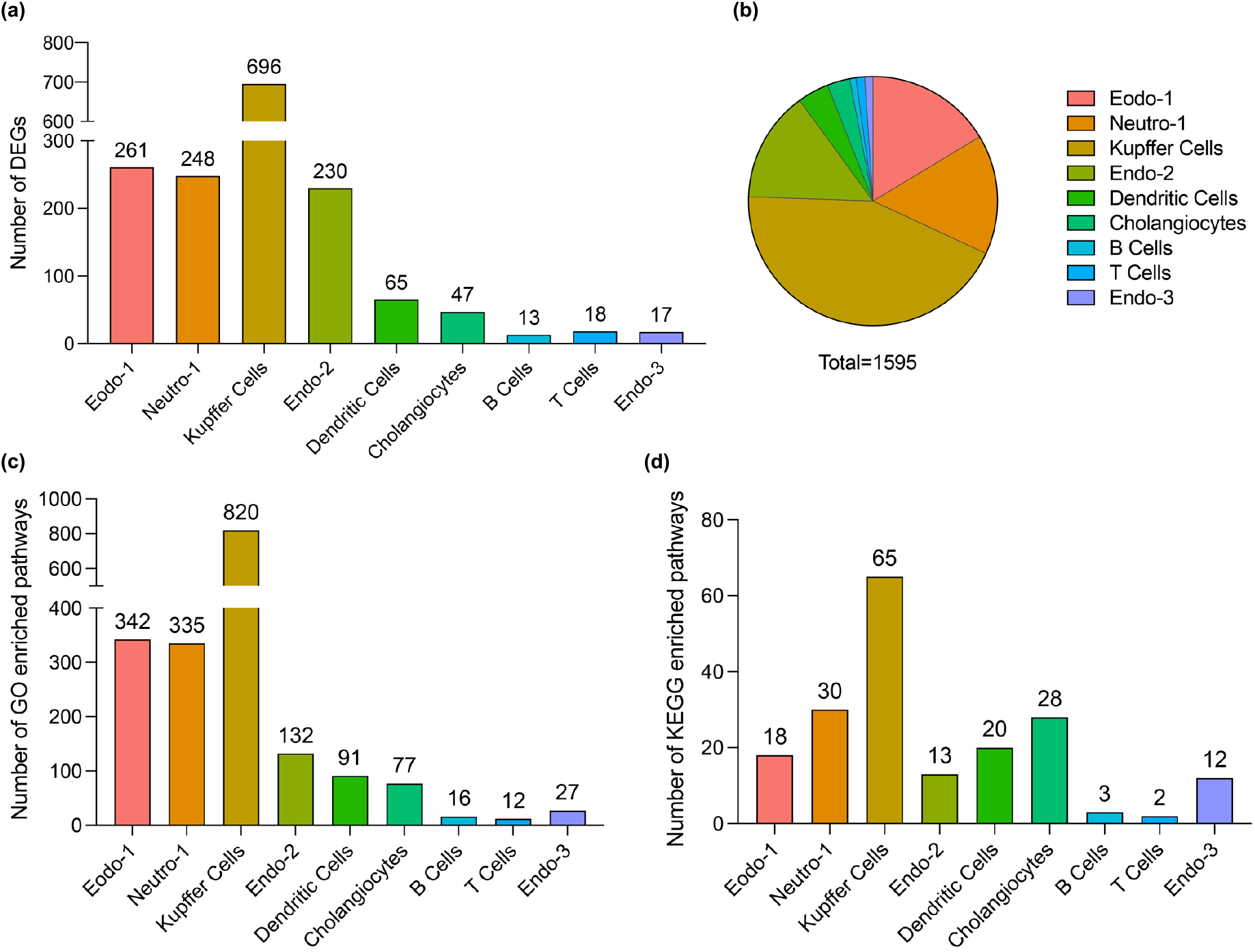
Cell-type-specific gene expression profiles and genetic pathways caused by PCB126. **(a)** DEGs between two groups of each cluster and **(b)** their proportion in total DEGs. **(c)** The number of GO biological process term enriched pathways. **(d)** The number of KEGG enriched pathways. Endo, endothelial cells; Neutro, neutrophils; DEGs, differentially expressed genes

By comparing the dataset against GO (biological process (BP)) and KEGG, the lists of GO BP enriched pathways and KEGG enriched pathways were obtained (**Fig. 2c,d**). It was discovered that different clusters had a different number of enriched pathways. For example, 696, 261,248 and 230 GO enriched pathways were found in Kupffer cells, Endo-1, Neutro-1 and Endo-2, respectively (**Fig. 2c**). Kupffer cells, Neutro-1 and cholangiocytes owned 65, 30 and 28 KEGG enriched pathways, respectively (**Fig. 2d**). Among all clusters, Kupffer cells were salient and possessed a much bigger number of enriched pathways than others.

### PCB126 induces the canonical genomic AhR pathway mainly in endothelial cells

To investigate the induction of the canonical genomic AhR pathway in different cell clusters, The gene expression profiles of *Ahr* (**Fig. 3a**) and *Arnt* (**Fig. 3b**) were inspected, respectively. In both two groups, cells exhibiting a high expression of two genes are homogeneously distributed in most cells, inferring that only some cells were in charge of expressing *Ahr* and *Arnt*. Among endothelial cells, Endo-2 and Endo-3 expressed *Ahr* and *Arnt* more than Endo-1. Besides, compared with the control group, PCB126 resulted in an upregulation of *Ahr* and *Arnt* in Endo-1. These results indicated that PCB126 differently influenced three clusters of endothelial cells.

**Fig. 3.**
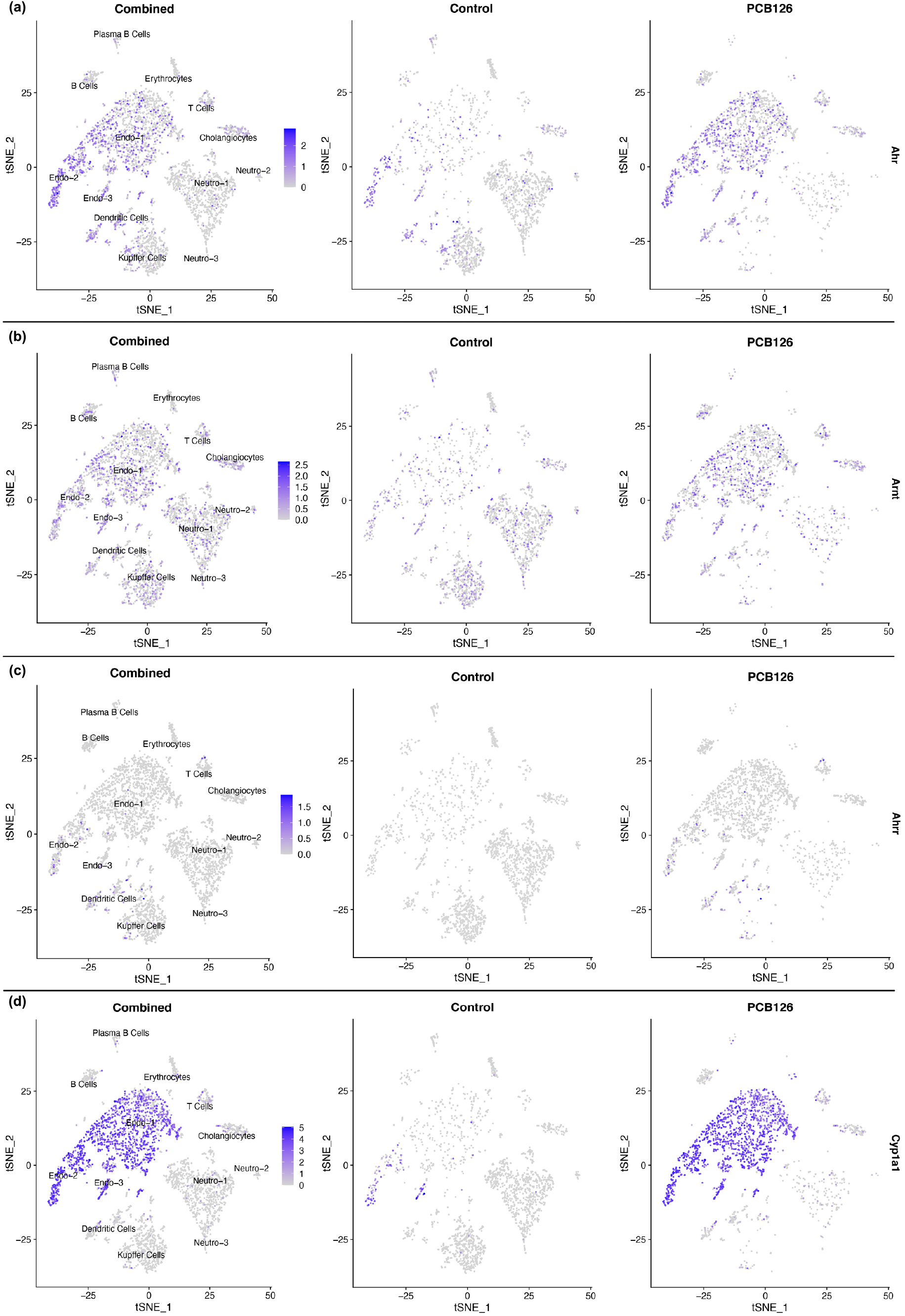
Gene expression profiles of **(a)** *Ahr*, **(b)** *Arnt*, **(c)** *Ahrr* and **(d)** *Cyp1a1* either combined or separated. Endo, endothelial cells; Neutro, neutrophils.

The gene expression profiles of AhR target genes like *Ahrr* and *Cyp1a1* were examined. All cells in the control group expressed no *Ahrr*, but in the PCB126 group, PCB126 induced a mild expression of *Ahrr* in some endothelial cells, dendritic cells, Kupffer cells and T cells (**Fig. 3c**). In terms of *Cyp1a1*, although Endo-2 and Endo-3 in the control group had a fraction of cells expressing *Cyp1a1*, this gene was significantly upregulated in all subpopulations of endothelial cells because of PCB126 exposure (**Fig. 3d**). These denoted that among all NPCs, PCB126 induced the canonical genomic AhR pathway mainly in endothelial cells. In addition to *Ahrr* and *Cyp1a1*, *Cyp1a2* and *Cyp1b1* which could be intrigued too (Manikandan and Nagini 2017) were also investigated. Between two groups, the gene expression profiles of *Cyp1a2* and *Cyp1b1* showed no significant difference (**Fig. S2**).

### PCB126 gives rise to immune responses in multiple immune cells

GO BP enriched pathways of immune cells were properly grouped into several themes by the software AutoAnnotate in Cytoscape. In B cells, enriched pathways were related to peptide process biosynthesis, cellular detoxification response, protein nitrosylation peptidyl, and neutrophil aggregation (**Fig. 4a**). Within cellular detoxification response, pathways were mainly associated with detoxification and oxidative stress.

**Fig. 4.**
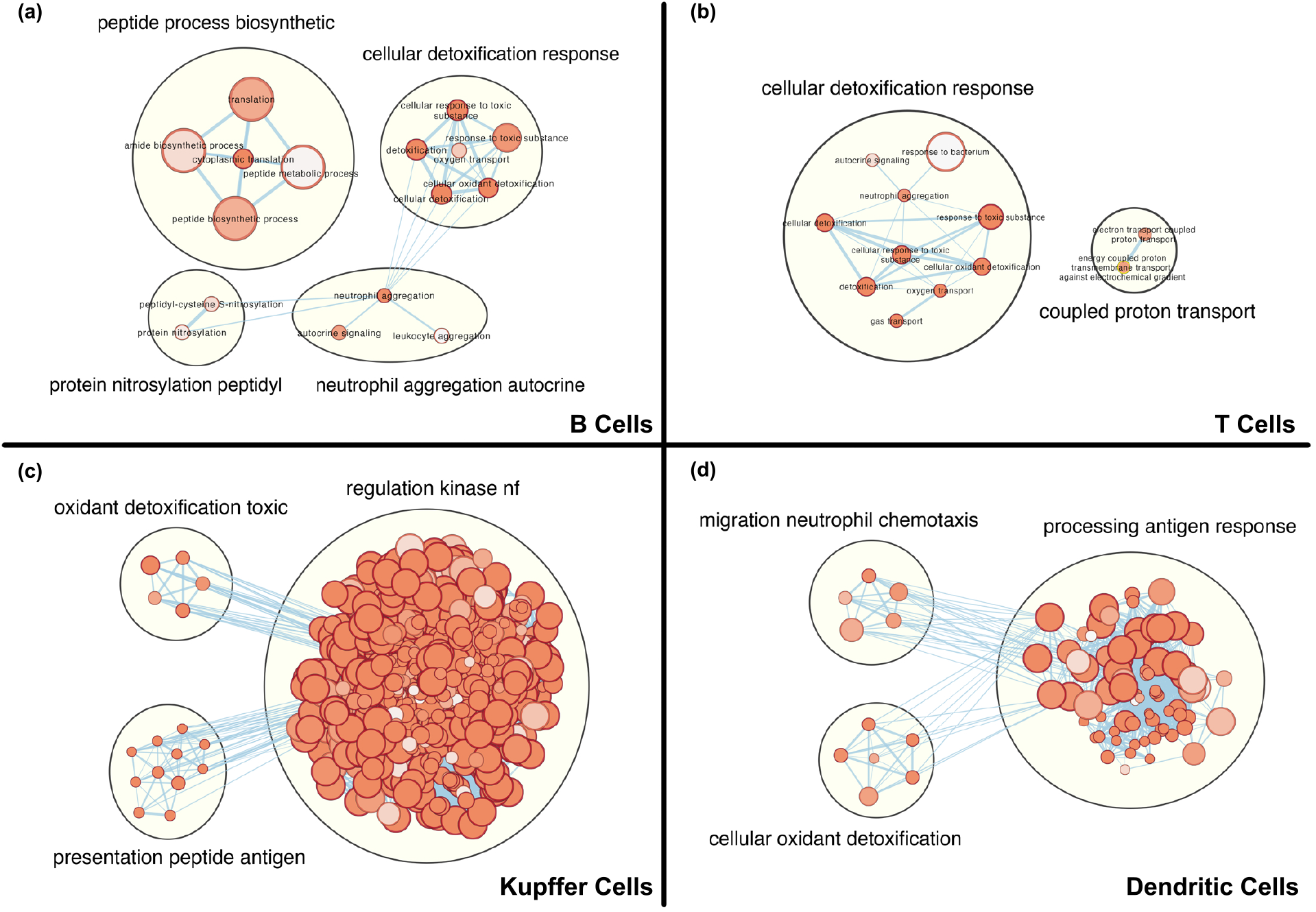
Main grouped themes of GO biological progress enriched pathways for **(a)** B cells, **(b)** T cells, **(c)** Kupffer cells and **(d)** dendritic cells.

In T cells, enriched pathways were divided into cellular detoxification and coupled proton transport (**Fig. 4b**). Most pathways were related to cellular detoxification response. Similar to the situation in B cells, enriched pathways in this group were mostly linked to detoxification and oxidative stress.

In Kupffer cells, PCB126 induced oxidant detoxification, presentation of peptide antigen, and regulation of kinase NF-κB (**Fig. 4c**). The transcription factor NF-κB plays a crucial role in the immune system. A well-known function of NF-κB is the regulation of inflammatory responses (Liu et al. 2017). Notably, most enriched pathways were related to the regulation of the kinase NF-κB. This denoted that Kupffer cells, the resident macrophages in the liver, were fighting against the inflammation, which may be caused by the presence of PCB126.

In the healthy liver, resident dendritic cells were mainly immature cells, inclined to capture and process antigens (Racanelli and Rehermann 2006). Once activated, they will downregulate antigen-related receptors and increase their ability to migrate (Matsuno et al. 1996; Kudo et al. 1997). In dendritic cells, PCB126 caused enriched pathways classified into migration of neutrophil chemotaxis, cellular oxidant detoxification, and processing antigen response (**Fig 4d**). Dendritic cells also showed signs of detoxification and oxidative stress. Apart from this, most pathways were related to processing antigen response, and their mobility was altered. These meant that dendritic cells were activated by PCB126. In summary, PCB126 significantly changed the transcriptional patterns of immune cells. It induced oxidative stress and detoxification in most immune cells, caused Kupffer cells’ response to inflammation and activated dendritic cells.

## Discussion

With a complicated architecture, the liver composes of various kinds of cells (Baratta et al. 2009). This pattern corresponds with its crucial immune and detoxification functions. Lots of scRNA-seq studies focusing on hepatic cells has shown this complexity (Halpern et al. 2017; MacParland et al. 2018; Aizarani et al. 2019). Regarding PCB toxicity, studies are often conducted on a bulk scale (e.g., bulk RNA sequencing (RNA-seq)), resulting in a loss of detailed information from a single-cell level. Apart from this, hepatic NPCs’ role in defending PCB126 tends to be marginalized and remains elusive. In this study, we applied scRNA-seq to resolve the toxic effects of PCB126 on hepatic NPCs with a single-cell resolution. By analyzing annotated cell clusters, identified DEGs and enriched pathways, we represented a detailed landscape of NPCs’ intricate responses to PCB126 exposure.

Recently, hepatic NPCs’ role as master regulators has been brought into focus. A cell-type-resolved proteome has revealed the division of labor between PCs and NPCs. It follows a system in which the former makes the main components of pathways, while the latter triggers the pathways (Ding et al. 2016). Besides, another similar study also indicates that NPCs may play a modulating role in the metabolism of xenobiotics and drugs (Ölander et al. 2020). It is proposed that hepatic NPCs play important roles in the pathogenesis of the alcoholic liver disease (Seo and Jeong 2016).

When handling xenobiotics like PCBs, they may act as pivotal regulators, too. In this study, we found that to fight against PCB126, NPCs played the role as regulators in their unique ways. Endothelial cells directly took part in clearing PCB126 through the activation of the canonical AhR genomic pathway. Immune cells’ transcriptional patterns were also altered. B cells, T cells, Kupffer cells, and dendritic cells all showed signs of oxidative detoxification. Kupffer cells handled inflammation through the regulation of kinase NF-κB. Resident dendritic cells were activated by changing their mobility and ability to capture antigens. In the presence of PCB126, hepatocytes hold the main task of PCB126 metabolism and excretion, but NPCs response and contribute, too. Interestingly, endothelial cells joined in the direct clearance of PCB126. In this case, they reacted just like hepatocytes. With the help of scRNA-seq, our study showed a whole picture of NPCs’ behaviors. This makes targeting specific cell type’ responses to PCB126 in an *in vivo* model possible. These findings are helpful since PCBs are still ubiquitous. In some places, workers or even residents are inevitably exposed to PCBs, although at a low level (Zhu et al. 2022). Dissecting cells’ responses to PCBs from a single-cell level helps identify remarkable cell types and pivotal altered pathways. By designing drugs or nutrients, activating, enhancing or inhibiting specific pathways seems to be promising in ameliorating the damage to human health caused by PCBs.

In addition to the difference in cells among different cell types, cells of the same type responded to PCB126 exposure differently. Endothelial cells in our dataset were divided into 3 subpopulations, which responded to PCB126 in a diverse manner. Many other scRNA-seq studies also discovered distinct subpopulations of hepatic endothelial cells (Halpern et al. 2017; MacParland et al. 2018). The liver consists of the basic building blocks called lobule. Along the lobule axis, blood flows from the portal vein to the draining central vein, forming the gradients of substances like nutrients, oxygen and toxins. This results in different gene expression patterns along the axis, a phenomenon termed zonation (Jungermann and Kietzmann 1996). This in part explains why cells of the same type (e.g., endothelial cells) can have different responses to PCB126, since they may actually contact different concentrations of PCB126 or its metabolites. Indeed, it was reported that PCB126 induced the activation of *cyp1a1* in vascular endothelial cells (Ramadass et al. 2003; Han et al. 2012). This is consistent with our findings. However, *in vitro* models or bulk RNA-seq could miss out on underlying information because of unbiasedness. The disparity among cells of the same type should not be ignored. It has already been reported that the average expression level of a cell population could be misleading and strongly biased, because a small fraction of cells with high or low expression in this population may blur the true picture (Bengtsson et al. 2005). ScRNA-seq empowers us to perform single-cell whole-transcriptome analysis (Wu et al. 2014) in an *in vivo* model. A transcriptomic and metabolomic analysis of HepaRG liver cells exposed to PCB126 showed several hallmarks of activation of the AhR receptor by dioxin-like compounds (Mesnage et al. 2018). In this study, by utilizing scRNA-seq, we went deeper and the canonical AhR pathway was delineated on a more explicit scale. That’s to say, in a single-cell level.

In our dataset, it was observed that in all cells, the transcriptional expression of *ahr* and *arnt* seemed to be quite interesting. Only part cells expressed two genes while the rest showed little sign of expression as if cells were able to communicate, cooperate and coordinate with each other. AhR has been detected in the serum (Hu et al. 2020). Producing target products in part cells and delivering them to all cells is a smart strategy since it is more energy-saving and space-saving. These gene expression patterns would certainly be ignored if bulk RNA-seq were used. Besides, the heterogeneity among cells described before will not be taken into consideration in a bulk RNA-seq study, either. However, it should be noted that we still don’t know the exact distribution of the protein AhR in every single cell. This challenge may be resolved in the future when single-cell proteomics (Marx 2019) comes into reality. An integration of single-cell transcriptome and single-cell proteome will be able to show a more comprehensive picture. From the scope of “precision toxicology”, many factors related to cell heterogeneity can be taken into consideration by using sophisticated technologies like scRNA-seq.

## Conclusions

Nowadays, as ubiquitous environmental pollutants, PCBs still pose great health risks. Here, by using scRNA-seq, we revealed hepatic NPCs’ intricate responses to PCB126 exposure in a mouse model. Cell-type-specific gene expressions and genetic pathways were unraveled. Besides, we also discovered activation of the AhR pathway, and immune responses more precisely. These results indicate that the heterogeneity among cells in different cell types, or most importantly, cells of the same type, should not be marginalized. Taken together, this study profiles PCB126 toxicity from a single-cell level and provides illuminating insights into the application of scRNA-seq in environmental toxicology.

