## Supplemental Information for "Single-cell transcriptomic analysis of mouse liver reveals nonparenchymal cells’ intricate responses to PCB126 exposure"

Address: Zhejiang University Zijingang Campus Environmental and Resource College Block B Room 327, No.866 Yuhangtang Road, Hangzhou 310058, Zhejiang China

This file includes 4 pages, 2 figures and 1 table.

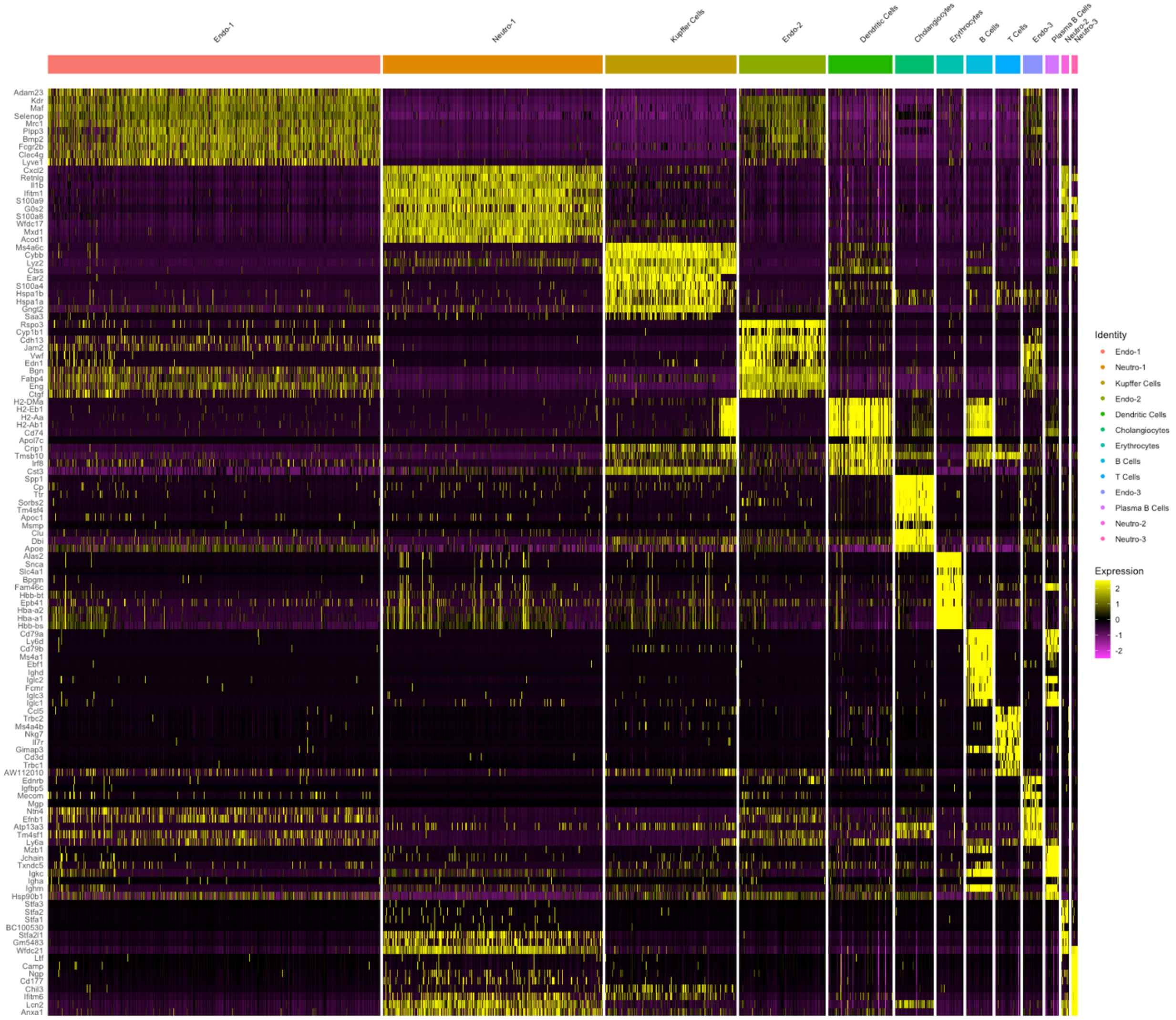


**Fig. S1** The heatmap of top 10 marker genes of each cluster. Top 10 marker genes were detected by the function FindAllMarkers of Seurat. Note: Endo, endothelial cells; Neutro, neutrophils

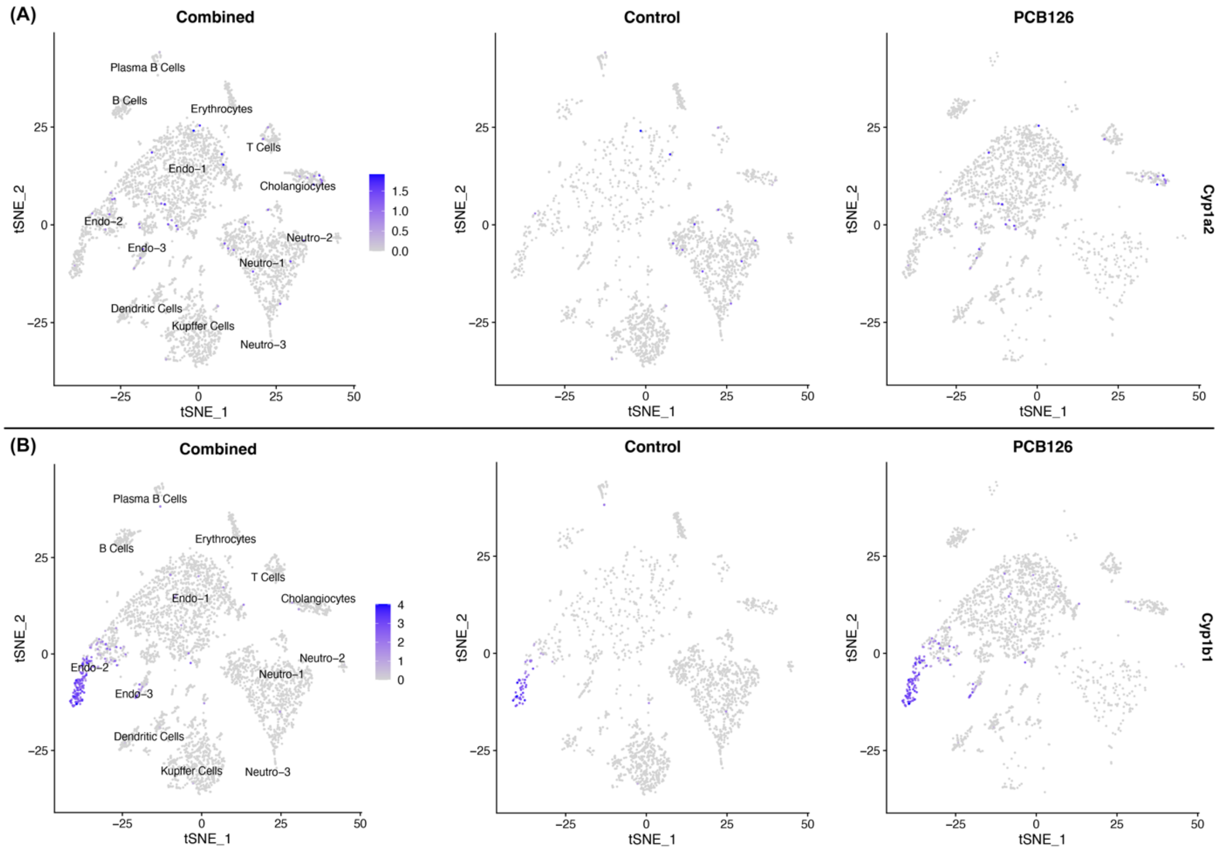


**Fig. S2** Gene expression profiles of **(a)** *Cyp1a2* and **(b)** *Cyp1b1* either combined or separated.

**Table S1** Cell types and their corresponding marker genes

| **Cell Types** | **Marker Genes** |
|---|---|
| **Endothelial Cells** | ***Clec4g, Kdr, Aqp1, Stab2*** |
| **Kupffer Cells** | ***Csf1r, Cd68, Ms4a6c, S100a4*** |
| **Neutrophils** | ***Csf3r, Lcn2, S100a8, S100a9*** |
| **Dendritic Cells** | ***Irf8*** |
| **Plasma B Cells** | ***Jchain, Ighm, Igkc, Cd79b*** |
| **B Cells** | ***Ebf1, Ighm, Igkc, Cd79b*** |
| **T Cells** | ***Cd3g, Trac, Trbc2, Trbc1*** |
| **Cholangiocytes** | ***Krt19, Epcam, Sox9*** |
| **Erythrocytes** | ***Epb41, Hba-a2, Hba-a1*** |
